# Off-the-shelf allogeneic polyclonal CD38KO/CD38-CAR γδT cells for the treatment of T cell malignancies

**DOI:** 10.1101/2025.08.30.673295

**Authors:** Genesis Snyder, Alexia K. Martin, Yasemin Sezgin, Colin Maguire, Branden S Moriarity, Beau R Webber, Erin Cross, Marcelo S. F. Pereira, Noushin Saljoughian, Ana L Portillo, Misaal Mehboob, Justin Lyberger, Kevin A Cassady, Ali A Ashkar, Gregory Behbehani, Dean A. Lee, Meisam Naeimi Kararoudi

## Abstract

Relapsed and refractory T cell malignancies are associated with poor clinical outcomes. Autologous sources of αβT cells have been employed for chimeric antigen receptor (CAR) therapies to eliminate the potential for graft-vs-host disease (GvHD). However, the application of CAR-T therapy for T-ALL has been hindered by an inability to obtain sufficient healthy αβT cells from patients combined with fratricide due to concurrent antigen expression on normal T cells. Here, we genetically engineered polyclonal γδT cells, which do not cause GvHD, as an allogeneic source for cancer immunotherapy targeting the pancancer antigen CD38. Utilizing a novel expansion protocol in combination with CRISPR/AAV gene editing, we developed CD38KO/CD38-CAR polyclonal γδT cells that target T-ALL. Our editing strategy enabled site-directed, on-target insertion of the CD38-CAR transgene into the CD38 locus, with no evidence of significant random CAR DNA integration (as commonly seen with lentiviral CAR transduction) or chromatin abnormalities resulting from CRISPR editing. This enhanced targeting effectively mitigated fratricide through simultaneous CD38 disruption and CAR expression. We demonstrated the efficacy of the CD38KO/CD38-CAR γδT cells in vitro across multiple patient-derived T-ALL samples collected at baseline and relapse. In vivo, a single injection of CD38KO/CD38-CAR γδT cells without exogenous cytokine support resulted in potent anti-leukemic efficacy. Fratricide-resistant CD38KO/CD38-CAR polyclonal γδT cells thus represent a promising off-the-shelf therapeutic platform for T cell malignancies and other CD38-expressing cancers.

**Key Points:**

- Hybrid pan-γδTCR antibody/mbIL21-41BBL feeder expansion yields high-purity, polyclonal γδT cells suitable for CRISPR/AAV editing.
- On-target CD38-CAR knock-in with simultaneous CD38 knockout prevents fratricide and enables potent T-ALL killing in vitro and in vivo.

## Introduction

Treating patients with relapsed and refractory T-cell acute lymphoblastic leukemia (T-ALL) remains a significant clinical challenge. While conventional therapies including allogeneic hematopoietic stem cell transplantation (HSCT) are widely used, long-term outcomes remain poor, with 5-year survival rates of approximately 25% in children and 50% in adults.^1-5^

Novel treatment strategies such as chimeric antigen receptor (CAR) T cells have shown remarkable success in malignancies expressing surface antigens like CD19, CD20, and BCMA.^6^ FDA-approved CAR-T cell therapies typically utilize autologous cells from the patient and are predominantly composed of alpha-beta (αβ) T cells. However, this approach is not optimal for T cell malignancies due to the high burden of malignant T cells within the patient’s own healthy T cell compartment.^3,4^ Additionally, most targetable antigens for T cell malignancies, such as CD70, CD7 and CD38, are shared between malignant and healthy T cells leading to fratricide during CAR-T cell manufacturing, thereby compromising yield and functionality.^1,3,7^

While the use of allogeneic donor-derived αβT cells is being explored including CD38CAR αβT cells,^8^ it necessitates genetic disruption of the TCRαβ/CD3 complex to prevent graftversus-host disease (GvHD), adding complexity and cost to CAR-T production.^9,10^ Alternatively, the use of natural killer (NK) cells as the base for CD38CAR cell therapy, which have been shown by us and others to be effective against CD38-expressing malignancies.^11,12^ However, NK cell products are likely to require multiple infusions due to the lower ability of NK cells to expand in vivo.^13^

Gamma-delta (γδ) T cells have garnered increasing interest in cancer immunotherapy owing to studies demonstrating their prognostic value in various malignancies.^14-16^ These cells act as a bridge between innate and adaptive immunity and can recognize transformed cells in a major histocompatibility complex (MHC)-independent manner.^16^ However, non-modified γδ T cells have so far shown limited clinical efficacy when used as an adoptive cell therapy.^14,17-19^ Engineering γδT cells with CARs has been proposed to enhance their antitumor potential.^14,20,21^ Lentiviral-based CAR transduction of γδT cells has proven technically challenging and is typically restricted to specific subtypes such as Vδ1 or Vδ2 due to their preferential expansion.^20,22,23^ Broader inclusion of Vδ subtypes is desirable given the evidence that different γδT cell subpopulations play crucial roles in immune surveillance across various tissues.^16,24^

To address this, we developed a method enabling robust expansion of diverse γδT cell subsets, including Vδ1+, Vδ2+, and Vδ1−/Vδ2− populations. We used a hybrid system, initiating with a short-term pan γδTCR antibody-based stimulation followed by expansion with mbIL21- and 41BBL-expressing feeder cells (FC). Building upon our previously reported isatuximab-based CD38-targeting CAR, which was successfully implemented in αβT and NK cells using a simultaneous CD38 knockout (KO) and CAR knock-in strategy to avoid fratricide, we now apply a similar genome editing approach to γδT cells.^11^ This avoids the cost of additional *TRAC*^*KO*^ required in CAR αβT cells to avoid GvHD and the high number of infusions needed with NK cells. To enable site-specific insertion of the CD38-CAR construct, we utilized CRISPR/AAV-mediated knock-in at the endogenous CD38 locus in expanded polyclonal γδT cells. Unlike lentiviral vectors which have low integration efficiency in γδT cells,^25,26^ our targeted editing approach resulted in a high rate of precise on-target insertion and no significant chromosomal abnormalities compared to non-edited γδT cells. This strategy also disrupts native CD38 expression, minimizing fratricide and enhancing therapeutic potential. Functionally, CD38KO/CD38-CAR γδT cells effectively eliminated CD38-expressing T-ALL cells in vitro and controlled disease progression in vivo. Taken together, these findings support CD38KO/CD38-CAR γδT cells as a promising immunotherapy that may facilitate a programmed allogeneic HSCT approach to induce molecular remission in relapsed/refractory T-ALL and other CD38+ malignancies.

## Methods

### γδT isolation and expansion

Human primary γδT cells were isolated from peripheral blood mononuclear cells (PBMCs) obtained from healthy donor buffy coats using density gradient centrifugation,^11,27^ followed by isolation by negative selection with the EasySep™ Human γδT Cell Isolation Kit (STEMCELL Technologies). Detailed information on study reagents is provided (Supplementary Table 1). The isolated γδT cells were expanded using two systems: the feeder cell (FC)-only system and the hybrid system.

In the FC-only system, freshly isolated γδT cells were stimulated by co-culture with genetically engineered K562 cells expressing mbIL21 and 41BBL at a 2:1 ratio on days 0, 7 and 14.^28^ On day 14, the cells were used for CAR-T production as described below.

In the hybrid system, isolated γδT cells were stimulated with 10 μg/mL plate-bound αPan-γδ TCR antibody (Miltenyi Biotec) and 2 μg/mL soluble αCD28 antibody (Tonbo Bioscience).^17^ For the first 2 days of stimulation, the cells were cultured in standard plates at a concentration of 1×10^6^ cells/mL, then transferred to G-Rex 6-well plates for an additional 5 days (total of 7 days). The cells were further expanded by FC stimulation for 7-10 days. Following expansion, the cells were used for CAR-T production as described below.

In all conditions, γδT cells were cultured in Human T Cell Media (Bio-Techne) supplemented with 5% CTS Immune Cell Serum Replacement (Gibco), antibiotics (penicillin–streptomycin), and 100 IU/mL human recombinant IL-2 which was replenished every 2–3 days. For expansion, 2.5×10^6^ γδT cells were co-cultured with 2.5-5×10^6^ FC cells at a beginning density of 0.5×10^6^ cells/cm^2^ in 6-well G-Rex culture plates (Wilson Wolf) for 7-10 days per stimulation cycle in a total volume of 100ml. Freezing was performed at the end of expansion periods (day 14 and 21 for FC-only system; day 14 or 17 for hybrid system) using CryoStor CS10 (STEMCELL Technologies). The purity of the isolated and expanded γδT cells was confirmed by flow cytometry (Supplementary Table 2).

### Generation of CAR-γδT Cells Using CRISPR and AAV

Cells were washed thrice with phosphate-buffered saline (PBS), pelleted, and resuspended in Lonza P3 buffer at a concentration of 3×10^6^ cells in 20 µl for small-scale electroporation, or 1.5×10^7^ cells in 100 µl for large-scale electroporation. The Cas9/RNP complex was prepared as previously described.^29^ In brief, HiFi CRISPR-associated protein 9 (Cas9) Nuclease V3 (Integrated DNA Technologies) was combined with guide RNA targeting exon 1 of the CD38 gene (5′-CTGAACTCGCAGTTGGCCAT-3′).^11,29,30^ Cells were electroporated using program EN-138 on a Lonza 4D nucleofector. Electroporated cells were then rested in culture medium for 30 minutes then transduced with a multiplicity of infection (MOI) of 75,000 or 150,000 of adeno-associated virus type 6 (AAV6; Andelyn Biosciences) carrying the CD38 CAR expression cassette flanked by CD38 homology arms, as previously described.^11^

The electroporated and transduced cells were rested in complete media in a 24-well plate (small scale) or 6-well plate (large scale) at a concentration of 1×10^6^ cells/ml. 24 hours post-transduction, the medium volume was doubled. On day 2 or 3 post-transduction, the cells were transferred to a 6-well G-Rex flask seeded with 3×10^6^ feeder cells (small scale) or at 2:1 ratio (large scale) in 100 ml medium supplemented with IL-2. A minimum of two additional expansions with FC were performed every 7 (FC-only) or 7-10 (hybrid) days, as described above.

### Cytogenetic analysis

Cytogenetic analyses using the KROMASURE™ K-Band, SCREEN, and InSite assays were performed on γδ T cells prepared at Nationwide Children’s Hospital. For SCREEN and InSite, cultures were incubated with DNA analogs from the dGH Cell Prep Kit (KROMATID, Cat # dGH 0001), arrested in first mitosis by a four-hour Colcemid block (KROMATID, Cat # COL 001), and fixed in freshly prepared 3:1 methanol:acetic acid. K-Band samples were processed identically without DNA analogs. All fixed samples were shipped to KROMATID for metaphase preparation.

K-Band assays followed standard G-banding (trypsin–Giemsa) to generate characteristic banding patterns for detection of structural and numerical abnormalities. Chromosome analysis adhered to ISCN guidelines.

The custom InSite assay was designed to assess CD38CAR integration and structural variation at the Chr4 locus. Probes included an ATTO-643–labeled CD38CAR probe (3.5 kb), an ATTO-550–labeled telomeric probe (Chr4: 14,809,769–15,768,219, GRCh38), and a 6-FAM–labeled centromeric probe (Chr4: 15,863,887–16,863,163). Oligos were tiled across unique genomic regions, repetitive sequences masked, and probe specificity validated by BLAST. Control spreads confirmed probe localization and hybridization fidelity. Experimental samples (wildtype and CD38KO/CD38CARv3 γδ T cells) were analyzed for on-target versus off-target integration and structural rearrangements including translocations, inversions, deletions, and insertions.

For whole-genome evaluation, SCREEN employed chromosome-specific fluorescent oligos hybridized to metaphase spreads, counterstained with DAPI, and imaged on an Applied Spectral Imaging Harmony system with a 100× objective. Karyograms from 50 cells enabled per-chromosome resolution of structural variation, providing ∼20 kb sensitivity for inversions, insertions, and translocations. Additional details on experimental protocols can be found in the Supplemental Methods.

### Metabolic assays

The extracellular acidification rate (ECAR) and oxygen consumption rate (OCR) of γδT cells were assessed with the Seahorse XFe96 Extracellular Flux Analyzer. To measure ECAR, γδT cells were washed and resuspended in Seahorse XF RPMI medium containing 2 mM L-Glutamine. To measure OCR, γδT cells were washed and resuspended in Seahorse XF RPMI medium containing 11.1 mM glucose, 1 mM sodium pyruvate, and 2 mM L-glutamine. γδT cells were plated at 2×10^5^ cells per well in at least triplicates on Poly-L-Lysine (Sigma Aldrich) coated XF96 culture microplates. ECAR was measured after sequential addition of 11.1 mM Glucose, 2 μM oligomycin, and 50 mM of 2-deoxy-D-glucose. OCR was measured after sequential addition of 2 μM oligomycin, 0.5 μM carbonyl cyanide p-trifluoro methoxyphenyl hydrazone (FCCP), and 0.5 μM rotenone/antimycin A. Extracellular expression of nutrient receptors were assessed by flow cytometry (Supplementary table 2).

### Mass cytometry assay for purity and lysis assay

For long-term lysis assays, T-ALL (Jurkat) cells were cultured in 6-well plates (precoated with rat tail collagen [Corning]) in serum-free medium. After 24 hours, WT or CD38KO/CD38CAR γδT cells were added at a ratio of 1:1 and co-cultured for 24 and 48 hours. Cells were stained and acquired on the Helios-CyTOF system as published previously (detailed antibody information can be found in Supplementary Table 3).^30,31^ SPADE clustering and analyses were performed as previously described.^30,32^

### Cytotoxicity assay on primary samples

Three-hour calcein AM (Invitrogen) cytotoxicity assays were performed on primary samples as previously described.^11^ In brief, frozen primary samples (baseline and relapse) were acquired from The Ohio State University Comprehensive Cancer Center Hematology Tissue Bank. The samples were thawed, immediately stained with calcein AM (4µg/mL), and co-cultured with effector cells at the indicated effector-to-target (E:T) ratios. Culture supernatant was collected after three hours and assessed by fluorimetry. Percent specific lysis was then calculated using Triton X-100 4% for the maximum control and untreated tumor cells for the spontaneous control. Cytotoxicity is plotted as percent specific lysis.

### In vivo studies

The Jurkat cell line was transduced with a luciferase expressing lentiviral vector. 8-to 10-week-old female NSG mice (Jackson Laboratory) were inoculated by intravenous (tail vein) injection with 10×10^6^ Jurkat cells in 200 µl of PBS. Mice were randomized and treated 3 days later with non-edited γδT cells (WT, 10×10^6^), CAR γδT cells (10×10^6^), or PBS (n≥5/cohort) in 200 µl of PBS by tail vein injection. Tumor engraftment and burden were monitored by bioluminescent imaging (IVIS Spectrum imaging system) following injection of D-luciferin.^3^ Mice were monitored for weight loss (20% cut-off), decreased activity, or hindlimb paralysis, and humanely euthanized when control cohorts met endpoint criteria. Blood, spleen, and bone marrow were collected for analysis by flow cytometry. Animal studies were performed in accordance with the Institutional Animal Care and Use Committee (IACUC) at Nationwide Children’s Hospital approved protocols (IACUC Protocol # AR24-00097).

## Results

### Expansion of γδT cells using hybrid pan-γδTCR stimulation followed by mbIL21/41BBL feeder cell co-culture preserves the polyclonal nature of the γδT cell and improves cell yield

Expansion of polyclonal γδT cells using either pan-γδTCR antibody stimulation or mbIL21/41BBL feeder cells (FC) have both been described before.^22,33^ However, the expansion rate of each remained modest. Therefore, we hypothesized that using a hybrid expansion system initiating with a short-term pan-γδTCR antibody stimulation followed by FC co-culture could induce γδT cells expansion (Figure 1A). Our results demonstrated that the hybrid system significantly improved cell yield within two expansions (∼10-fold, 7.8x10^7^ cells vs 6.9x10^6^ cells, lognormal *t*-test, *p* = 0.0053) compared to the FC-only expansion system (Figure 1B). We confirmed high purity of γδT cells (detected as CD3^+^αβ T^-^ cells) post-expansion with an increase in CD3^+^ cells and a decrease in CD3^-^ cells in both expansion systems by flow cytometry. There was a marked reduction in CD3^+^αβ T cells (from 64% in PBMCs to 0.3% in expanded γδT cells, paired *t*-test, *p* = 0.008), an increased frequency of CD3^+^αβ T^-^γδTCR^+^ γδT cells (from 3% to 82%, *p* = 0.009), and a reduction in CD3^−^CD56^+^ NK cells (from 19% to 1.8%, *p* = 0.02) (Figure 1C and 1D). Both expansion methods maintained subtype diversity over three weeks (Figure 1E). Thus, regardless of prior pan-γδTCR antibody activation, FCs supported the expansion of polyclonal γδ T cells. Mass cytometry (CyTOF) further confirmed product purity and preserved polyclonality in expanded γδT cells (Figure 1F). Importantly, no CD3^+^αβ T cells were detected, supporting potential clinical-grade manufacturing devoid of GvHD-inducing contaminants (Figure 1F).

**Figure 1.**
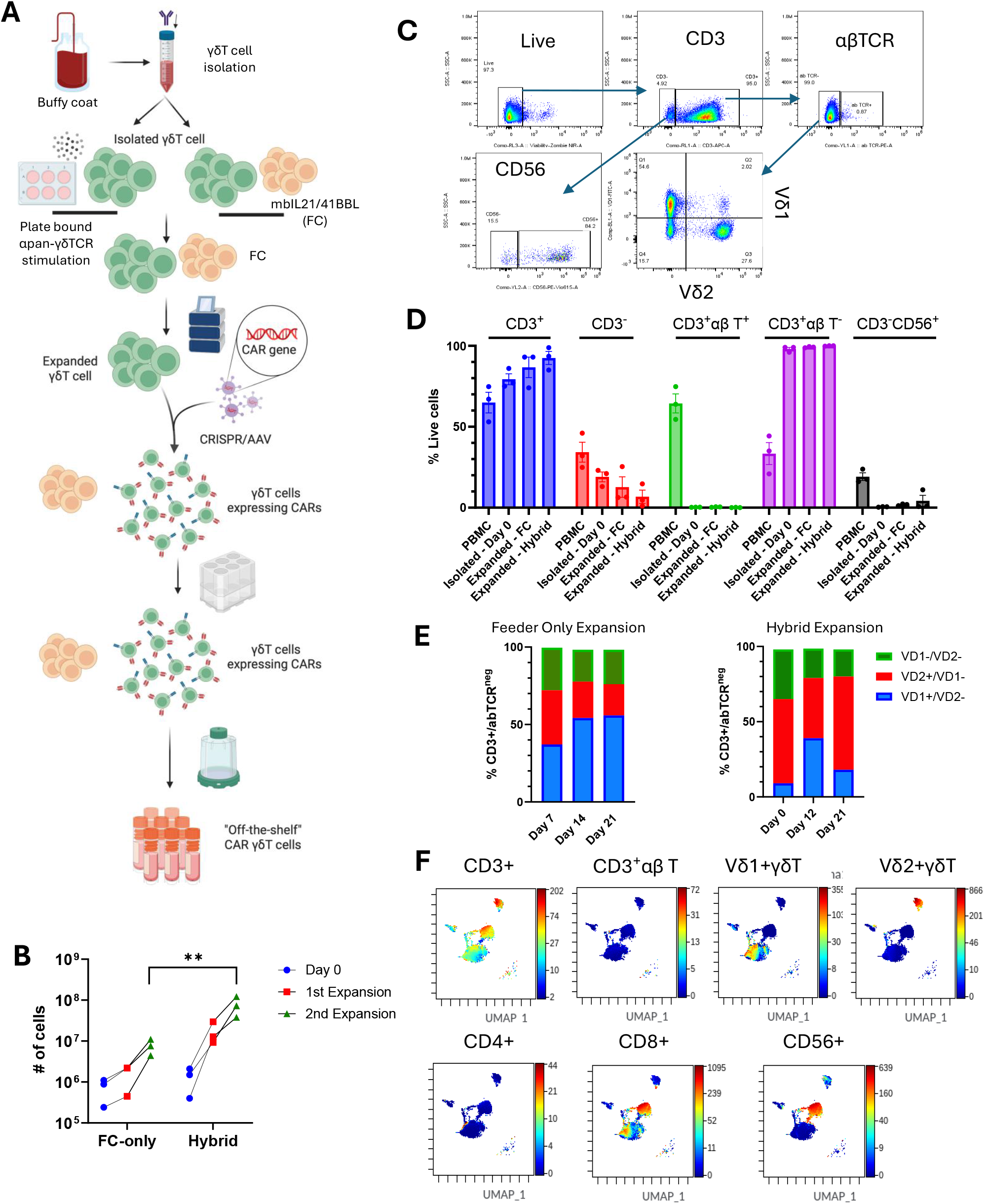
Expansion and Characterization of Polyclonal γδT Cells. (A) Schematic overview of γδT cell expansion protocols using hybrid system (αPan-γδ TCR antibody stimulation followed by feeder cell [FC] expansion) or FC-only expansion system, along with subsequent CAR γδT cell manufacturing and expansion steps. (B) Expansion of γδT cells using the FC-only and hybrid systems (change in γδT cell yield from day 0 to second expansion, lognormal *t*-test, *p* = 0.005). (C) Representative flow cytometry plots showing the gating strategy on the γδT cells to study their purity. (D) Flow cytometry analysis confirms high purity of γδT cells post-expansion, with minimal contamination from CD3^+^αβ T cells or CD3^−^CD56^+^ NK cells (mean ± standard error of the mean) and (E) both expansion platforms maintained γδT cell subtype diversity over a three-week culture period, preserving the polyclonal nature of the population (n = 3 healthy donors). (F) Mass cytometry (CyTOF) profiling revealed a preserved polyclonal composition and validated the immunophenotypic purity of the expanded γδT cell product. Color gradient of marker expression overlaid on Uniform Manifold Approximation and Projection (UMAP) plots of γδT cells.

### Generation of CRISPR/AAV-edited CD38KO/CD38CAR γδT cells

We next sought to generate CD38KO/CD38CAR γδT cells. Electroporation with CD38-targeting Cas9/RNP resulted in a substantial reduction in CD38 expression in CD38KO γδT cells relative to WT in the FC-only expansion system (9% vs 99%, *p* < 0.0001) and hybrid-expansion system (14% vs 71%, *p* < 0.0001) (Figure 2A and 2B). No significant difference in CD38 knock-out efficiency was observed between the two expansion systems. Following electroporation, γδT cells were transduced using two different multiplicities of infection (MOIs), 75K and 150K, of AAV6 to generate CD38KO/CD38CAR γδT cells. We used our previously reported isatuximab-based CD38CARv3 construct. CD38KO/CD38CAR γδT cells were generated at two different scales: a small scale (3×10^6^ cells) and a clinically relevant large scale (15×10^6^ cells). On average, CAR expression reached 38% with the hybrid system (WT vs CAR, *p* = 0.002) and 22% with the FC-only system (WT vs CAR, *p* = 0.03) (Figure 2C and 2D). Although not statically significant, CAR expression was higher in γδT cells expanded in the hybrid system. Importantly, there were no significant differences in CAR expression between the two scales or MOIs tested, highlighting the clinical feasibility of this protocol (Figure 2D).

**Figure 2.**
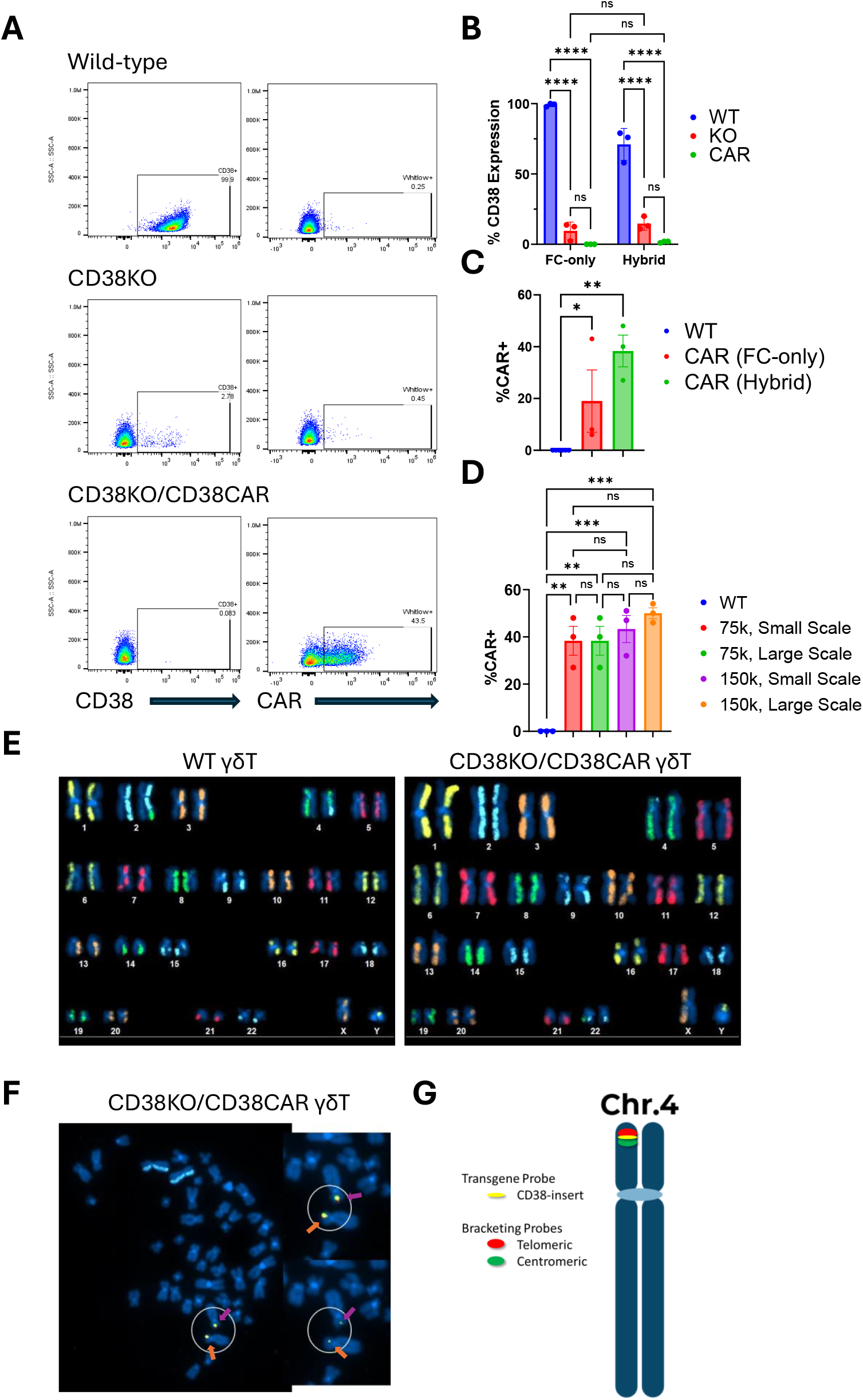
Successful generation of CD38KO/CD38CAR γδT Cells. (A) Representative flow cytometry plots showing the expression of CD38 and CD38CAR in γδT cells. (B) CRISPR-mediated targeting of CD38 in γδT cells significantly reduced CD38 expression, as assessed by flow cytometry in FC-only and hybrid expansion systems. Residual CD38^+^ γδT cells underwent fratricide, resulting in complete loss of detectable CD38 on CD38KO/CD38CAR γδT cells in FC-only and hybrid expansion systems. (C) The CRISPR/AAV approach achieved high surface expression of CD38CAR on γδT cells expanded using the FC-only or hybrid system. (D) Both small- and large-scale manufacturing of CD38CAR γδT cells yielded efficient CAR generation, with no significant differences in CAR expression between two multiplicities of infection (MOIs) of AAV6 (75,000 vs. 150,000). (E) Representative karyotype (K-band) analysis of wild-type and CD38KO/CD38CAR γδT cells. (F) InSite analysis confirmed precise, on-target integration of the CAR transgene, demonstrating biallelic integration. Two cells with CAR integration at both CD38 alleles are circled and identified by colored arrows (orange or pink) in each image. Enlarged images (right) display transgene signals in a separate channel (bottom right). (G) Integration was detected using probes specifically designed for the CD38CAR transgene and CD38 insertion site on chromosome 4, as shown in the schematic. Statistical analyses were performed by two-way ANOVA (B) or one-way ANOVA (C-D) with Holm-Šídák multiple comparison corrections. Data generated from 3 healthy donors and presented as the mean ± standard error of the mean. ns, not significant. \**p* < 0.05, \*\**p* < 0.01, \*\*\**p* < 0.001, \*\*\*\**p* < 0.0001.

### Cytogenetic analysis shows successful site directed insertion of CAR and no significant CRISPR/AAV mediated chromosomal abnormalities

We have previously shown the low off-target effect of the CD38 targeting gRNA used in this study.^27^ Here, we further investigated the off-target insertion and chromosomal abnormalities post CRISPR/AAV gene editing. Across all three assays - K-Band, SCREEN, and InSite - both the edited CD38KO/CD38CAR γδT and WT γδT-cell populations displayed overall diploid karyotypes with low levels of structural variation. K-Band analysis detected no recurrent abnormalities involving the CD38 target chromosome (Chr 4) in either sample. Aneuploidy was significantly lower in the edited sample compared to wildtype (5% vs. 17%, *p* = 0.011), driven entirely by fewer whole-chromosome losses; other structural variants occurred at low frequencies in both samples with no statistically significant differences.

SCREEN results were consistent with these findings, showing no significant differences in event rates for Chr 4, including no enrichment of inversions, SCEs or translocations at the target locus. Both samples exhibited a recurrent small inversion on Chr 8p in 100% of cells, homozygous in most, which is likely a germline variant in the donor. Inversions and SCEs were the most frequent event types, with the edited sample showing slightly fewer total events (301 vs. 317 in wildtype), slightly lower aneuploidy (8% vs. 10%), a single insertion not present in the wildtype, modestly more translocations (3 vs. 2 cells), and more complex events (4 vs. 1 cell), (Figure 2E).

InSite analysis confirmed on-target integration of the CAR transgene in 110 of 200 γδT cells, with 29 showing biallelic integration, and no structural rearrangements, translocations, truncations, or complex events involving the CD38 locus in either sample. Off-target insertions were observed in 44 CD38KO/CD38CAR γδT cells, appeared randomly distributed, were non-clonal, and did not exceed two integrations per cell, consistent with expectations for a safe drug product profile, particularly when contrasted with lentiviral approaches, where random integration is inherent (Figure 2F and 2G).

Together, these results indicate that the editing process did not introduce detectable clonal structural variation at the target locus and that the overall genomic integrity of the edited cells was comparable to the wildtype control.

### Effect of gene engineering on γδT cells metabolism

We next studied the effect of gene editing and the deletion of CD38, an NADase enzyme, on the metabolism of γδT cells. We measured the expression of nutrient receptors, extracellular acidification rate (ECAR), and oxygen consumption rate (OCR) of CD38KO γδT cells and CD38KO/CD38CAR γδT cells compared to WT γδT cells. Relative to WT controls, we observed no difference in the percent surface expression of Glut1 (glucose transporter 1) and CD71 (iron transporter), or MFI of CD98 (amino acid transporter) on γδT cells lacking CD38 expression (Supplementary Figure 1). No differences were observed in basal glycolysis, glycolytic capacity, or glycolytic reserve between CD38KO, CD38KO/CD38 CAR, and WT γδT cells. Similarly, CD38KO and WT γδT cells had comparable basal and maximal respiration. Lastly, spare respiratory capacity and mitochondrial linked-ATP production of CD38KO γδT cells were equivalent to WT γδT cells. Overall, these results demonstrate that gene editing does not affect the metabolic capacity of γδT cells. Although we had previously shown that CD38KO can enhance metabolism of NK and αβT cells,^11,27^ deletion of CD38 did not improve the glycolytic or oxidative metabolism of γδT cells.

### CD38KO/CD38CAR γδT cells exhibit potent antitumor activity in vitro

To evaluate the antitumor activity of CD38KO/CD38CAR γδT cells against T-ALL, we performed a long-term CyTOF-based cytotoxicity assay by co-culturing Jurkat cells (αβTCR^+^ T-ALL) with either WT or CD38KO/CD38CAR γδT cells for 24 and 48 hours. CyTOF analysis revealed a significant reduction in viability of the αβTCR^+^ Jurkat cells, indicated by phosphorylated retinoblastoma protein (pRB) expression, within 24 hours in the CAR γδT-treated group (*p* = 0.013) (Figure 3A and 3C). Although not statically significant, the lowest proportion of CD38+ Jurkat cells was observed in the CAR γδT-treated group at 48 hours post co-culture (Figure 3B and 3D). These findings demonstrate both the potency and specificity of CD38KO/CD38CAR γδT cells.

**Figure 3.**
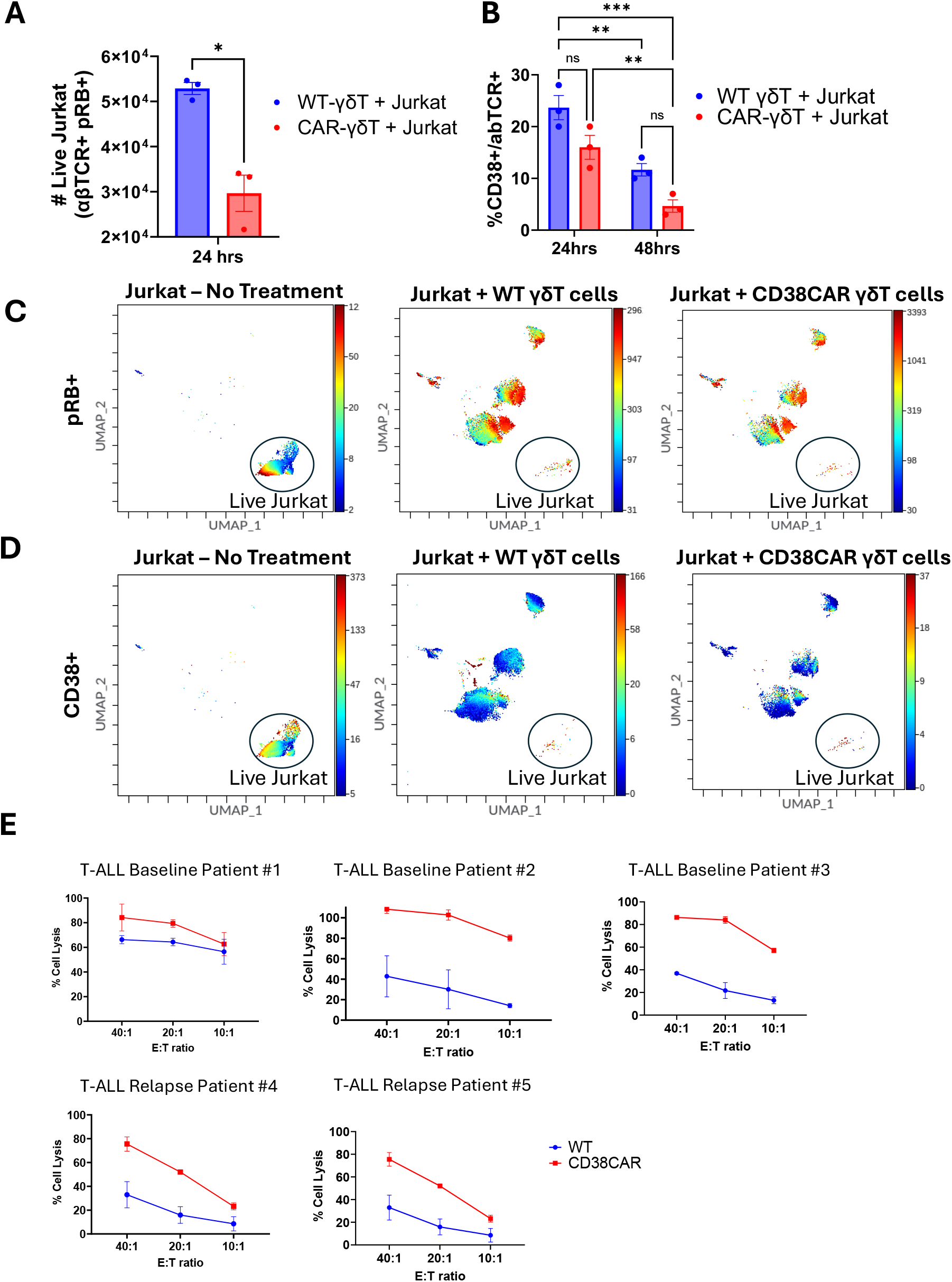
Antitumor activity of CD38KO/CD38CAR γδT cells against Jurkat and primary T-ALL samples. (A) Long-term CyTOF-based cytotoxicity assay in which Jurkat cells (αβTCR^+^ T-ALL) were co-cultured with either wild-type (WT) or CD38KO/CD38CAR (CAR) γδT cells for 24 hours. CAR γδT cells significantly reduced the number of live Jurkat cells, measured as pRB^+^ αβTCR^+^ cells (paired *t*-test, p = 0.013, n = 3 donors). (B) CAR γδT treatment also significantly decreased the number of CD38 expressing αβTCR^+^ Jurkat cells over time, confirming CAR specificity. Statistical analyses were performed using two-way ANOVA with Holm-Šídák multiple comparison correction (n = 3 donors). (C-D) Representative Uniform Manifold Approximation and Projection (UMAP) plots showing marked reduction of live (pRB+) CD38^+^ Jurkat cells following γδT cell treatment. (E) CD38KO/CD38CAR γδT cells exhibited enhanced antitumor activity against both baseline and relapsed primary T-ALL samples relative to WT γδT cells. For each assay, one donor-derived γδT cell (WT and CAR) was tested in triplicate. Data presented as mean ± standard error of the mean. ns, not significant. \**p* < 0.05, \*\**p* < 0.01, \*\*\**p* < 0.001.

To assess the clinical relevance of CD38KO/CD38CAR γδT cells, we evaluated their antitumor activity against primary T-ALL cells obtained from patients at both baseline and relapse stages using a 3-hour Calcein-based cytotoxicity assay. Despite the technical challenges of culturing thawed primary samples, we successfully performed the assay on three baseline and two relapsed patient samples using WT or CD38KO/CD38CAR γδT cells. Across all samples, CD38KO/CD38CAR γδT cells consistently exhibited superior anti–T-ALL activity compared with WT γδT cells across multiple effector-to-target (E:T) ratios (Figure 3E).

### CD38KO/CD38CAR γδT Cells Exhibit Potent Antitumor Activity Against T-ALL In Vivo

To assess the in vivo potency of CD38KO/CD38CAR γδT cells, we infused luciferase-expressing Jurkat T cells (Luc^+^ Jurkat) into NSG mice. Three days post-injection, tumor engraftment was confirmed via bioluminescent imaging and mice were treated intravenously with one of the following: non-edited, wild-type [WT] γδT cells, CAR γδT cells, or PBS (Figure 4A). Tumor progression was monitored weekly using bioluminescent imaging. By day 24, mice treated with PBS or WT γδT cells reached humane endpoints due to weight loss, consistent with previously reported findings using Jurkat xenografts (Figure 4B).^3^ Similarly, PBS and WT γδT treated mice had lost significantly more weight from baseline relative to CAR γδT cell treated mice (CAR γδT vs PBS, *p* = 0.0022; CAR γδT vs WT γδT, *p* = 0.0025). At this time, all mice were euthanized for analysis of tumor burden by flow cytometry. Ventral-side bioluminescence confirmed reduced tumor burden in CAR γδT-treated group vs PBS (*p* = 0.02) and WT γδT treated mice (*p* = 0.02) (Figure 4C-4D). A similar trend was observed in dorsal-side bioluminescence which demonstrated reduced tumor burden in the CAR γδT-treated group compared to WT γδT (*p* = 0.04) and PBS (*p* = 0.002) treated cohorts (Figure 4E). These imaging findings were further confirmed by flow cytometry, which revealed a substantial clearance of Jurkat cells (live CD3^+^/αβTCR^+^) from the spleen and bone marrow in the CAR γδT treated mice (Figure 4F-H). Relative to PBS-treated controls, mice treated with CAR γδT had a significantly lower proportion of CD3^+^/αβTCR^+^ cells in the spleen (*p* = 0.006) and bone marrow (*p* = 0.006). A similar trend was observed in peripheral blood, though not statistically significant (Supplemental Figure 2). Furthermore, of the limited remaining CD3^+^/αβTCR^+^ cells (2-18 cells) in samples from CAR γδT cell treated mice, the proportion expressing CD38 was significantly lower compared to PBS-treated mice in the spleen (*p* = 0.03) and bone marrow (*p* = 0.04). Taken together, these results demonstrate the anti-tumor efficacy of the CD38KO/CD38CAR γδT cells against T-ALL *in vivo*.

**Figure 4.**
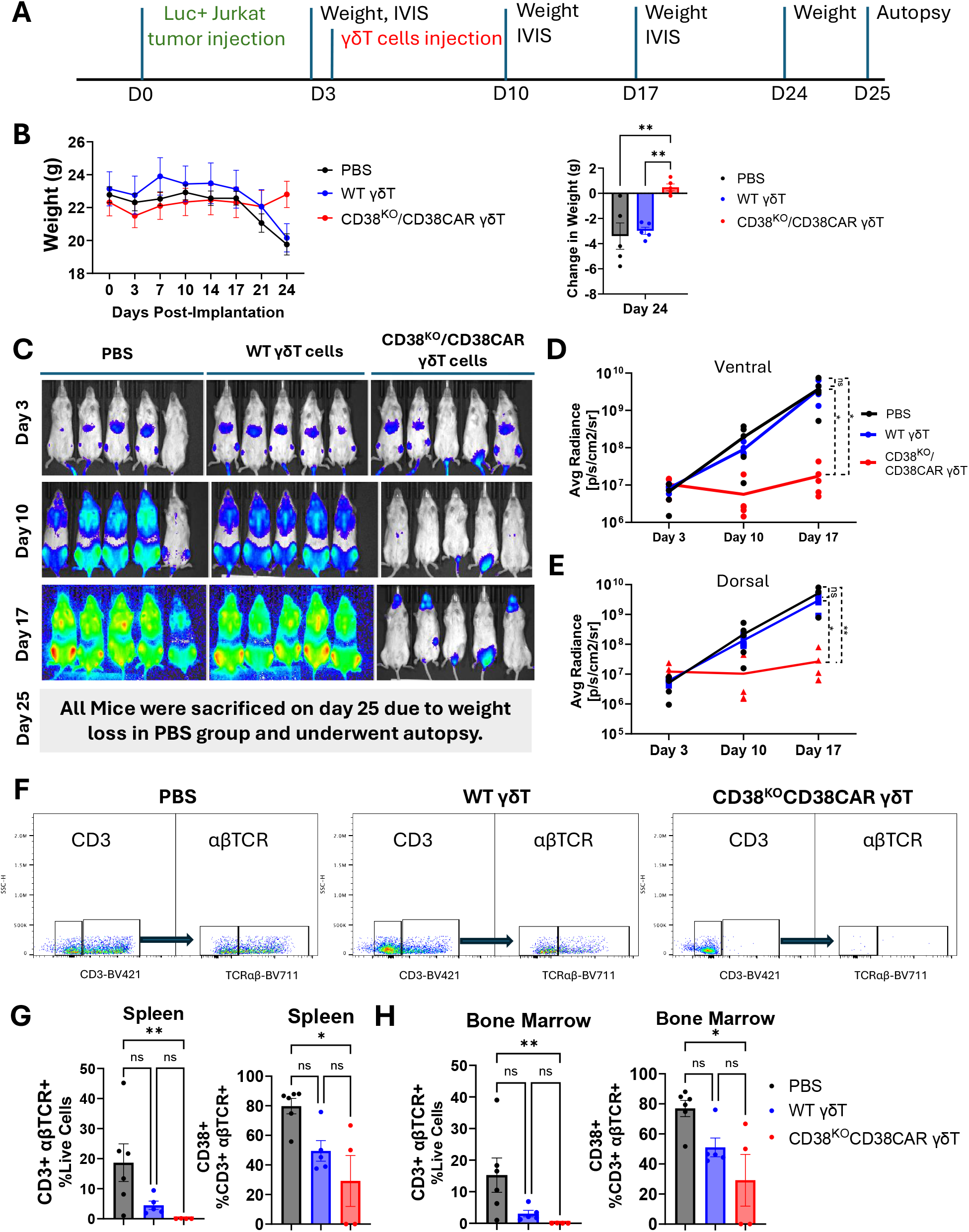
CD38KO/CD38CAR γδT Cells exhibit potent antitumor activity against T-ALL in vivo. (A) NSG mice engrafted with Luc^+^ Jurkat cells were treated with PBS, WT γδT, or CD38KO/CD38CAR γδT cells (n≥5 per cohort). (B) Mice treated with WT γδT cells or PBS had significantly greater weight loss from baseline (day 0) on day 24 relative to CD38KO/CD38CAR γδT cell treated mice (CAR vs WT γδT *p* = 0.0025; CAR vs PBS *p* = 0.0022). (C) Tumor burden estimated by bioluminescence imaging in PBS, WT γδT, and CD38KO/CD38CAR γδT treated cohorts. Tumor growth curves approximated by bioluminescent imaging (average radiance) of PBS, WT γδT, and CD38KO/CD38CAR γδT treated mice from ventral (D) and dorsal (E) side of the mice with differences in tumor burden analyzed on day 17. (F) Representative flow cytometry plots of CD3 and αβTCR expression. Tumor burden (CD3+ αβTCR^+^ T-ALL) and CD38+ expressing CD3+ αβTCR^+^ cells were measured in samples from spleen (G) and bone marrow (H) collected at autopsy. Statistical analyses were performed using one-way ANOVA with Holm-Šídák multiple comparison correction (B, D, E) or non-parametric Kruskal-Wallis test with Dunn’s multiple comparison correction (G, H). Data presented as mean ± standard error of the mean with each shape representing one animal. d, day; g, grams; avg, average; ns, not significant. \**p* < 0.05, \*\**p* < 0.01.

## Discussion

Current FDA-approved CAR-T cell therapies for relapsed/refractory cancers are derived from autologous αβT cells. However, this approach presents major limitations, particularly for patients with T cell malignancies. Key challenges include: 1. presence of malignant T cells (e.g., T-ALL) making isolation of healthy autologous T cells difficult for CAR-T manufacturing, 2. manufacturing delays, 3. high production costs of bespoke manufacturing (>$400,000/patient) limiting accessibility and affordability, and 4. fratricide caused by shared antigens leading to CAR-T cells attacking one another.^34,35^ Generating CAR-T cell products using allogeneic αβT cells also presents challenges including risk of severe graft-versus-host disease (GvHD) and fratricide due to shared antigens.^35,36^ These barriers highlight the urgent need for alternative T cell sources. γδT cells are a promising prospective T cell source with the potential to mitigate some of these challenges. However, their efficacy as an adoptive cell therapy has remained limited, suggesting a need for further engineering to enhance their antitumor activity.^14,17-21^

Although allogeneic CAR-T cells targeting other T-ALL antigens such as CD2, CD5, CD7, and CD70 have been developed and tested, CD38 is recognized as a pan-targeting antigen across multiple hematological malignancies, not just T-ALL.^2,4,37-39^ Importantly, CD38 expression is consistently high in T-ALL across different disease stages and subtypes, including precursor, thymic, and mature T-ALL.^40^ Moreover, CD38 expression is significantly higher on T-ALL cells than on normal T cells.^40,41^ It has been reported that CD38 is expressed on more than 80% of T-ALL cells at MRD (88.7%) and in relapsed cases (82.9%).^40^ We previously demonstrated that an isatuximab-based CD38CAR, inserted into the CD38 gene, can overcome fratricide in both αβT cells and NK cells.^11^ Thus, we sought to leverage this editing approach for engineering γδT cells. Our method enables robust expansion of all γδT cell subsets and establishes optimal conditions for gene editing and large-scale clinical manufacturing. Here, we describe a highly efficient method for genetic engineering of polyclonal γδT cells, including Vδ1^+^, Vδ2^+^, and Vδ1^−^/Vδ2^−^ subsets. As subtypes of γδT cells play distinct roles in responding to cancer and infection, preservation of polyclonality may be critical for anti-tumor efficacy.^16,24^ The prevalence of different subtypes across tissues may represent unique biological properties that contribute to controlling T-ALL in both peripheral blood and tissues. This was evident in our mouse model, where we observed a reduction of T-ALL burden in multiple tissue compartments (e.g., bone marrow, spleen, blood). Our technology represents a promising alternative cell therapy based on γδT cells’ inherent anti-tumor activity, lack of GvHD induction, suitability for off-the-shelf use, and in vivo persistence. Here, we demonstrate the efficacy of a novel CD38CAR γδT cell product that is effective against multiple T-ALL subtypes, with activity both in vitro and in vivo. These results establish the feasibility of using allogeneic polyclonal CD38KO/CD38CAR γδT cells as a potential treatment to reduce tumor burden and serve as a bridge to allogeneic HSCT. Beyond T-ALL, our off-the-shelf CD38CAR γδT cells also have potential therapeutic applications in other CD38^+^ malignancies.

## Acknowledgements

We would like to thank CancerFree KIDS Foundation, Wilson Wolf Manufacturing, and the Office of Technology Commercialization at Nationwide Children’s Hospital for providing funding for this project. A.K.M., K.A.C., D.A.L., G.B., and M.N.K are supported by NIH (U54-CA232561). A.K.M and K.A.C are supported by DOD (HT9425-23-1-0803). A.K.M is supported by NIH (T32-CA269052). Additionally, we would like to thank the Flow Cytometry Core, Animal Research Core, and the Preclinical Behavioral Imaging Core at Nationwide Children’s Hospital for their assistance conducting the *in vivo* study.

## Authorship

Authorship Contributions: G.S., A.K.M., Y.S., C.M., E.C., M.S.FP., N.S., A.L.P., M.M., and J.L. performed the experiments; B.S.M., B.R.W., E.C., K.A.C., A.A.A., D.A.L., and G.B. provided valuable reagents and intellectual input; A.K.M., G.S., Y.S., C.M., and M.N.K. designed the research and analyzed the data; A.K.M., G.S., E.C., and M.N.K. wrote the manuscript which was reviewed by all authors.

Disclosure of Conflict of Interest: D.A.L., M.N.K., G.S., N.S., A.K.M and Y.S. have a provisional patent for the described technology which has been licensed to CARTx Therapeutics. M.N.K. is the CSO and co-founder of CARTx Therapeutics. D.A.L., M.N.K., and M.S.F.P. have received royalties from Sanofi/Kiadis. B.M. and B.R.W. have submitted a patent for part of the technology described here. E.C. is a Kromatid employee. The remaining authors declare no competing financial interests.

