## Supplementary Material for "Off-the-shelf allogeneic polyclonal CD38KO/CD38-CAR γδT cells for the treatment of T cell malignancies"

**Supplementary Methods**

**Sample preparation for cytogenetic analysis**

Sample preparation for cytogenetic analysis with the KROMASURE™ K Band, SCREEN, and InSite assays was using γδT Cells generated from one donor. At the end of 10 days expansion with FC stimulation (culture conditions described above), $2.5\times10^6$ γδ T cells were plated in two T25 flask at $0.5\times10^6$cell/mL concentration, one flask designated for K Band and the other for SCREEN and InSite assays.

**For the SCREEN and InSite assays,** culture medium was replaced with a solution containing DNA analogs from the directional genomic hybridization (dGH) Cell Prep Kit (KROMATID, Cat # dGH 0001). Cells were arrested in first mitosis via a four hours Colcemid block (KROMATID, Cat # COL 001), harvested, and fixed in freshly prepared 3:1 methanol:acetic acid fixative (Fisher Scientific, Waltham, MA, USA). The K-Band samples were prepared identically, except that no DNA analogs were added. All fixed samples were then shipped to KROMATID for metaphase spread preparation. G-banded metaphases for K-Band were generated using the standard G-banding protocol, trypsin digestion followed by Giemsa staining, which produces characteristic dark (AT-rich) and light (GC-rich) bands that allow detection of structural and numerical chromosomal abnormalities (Howe et al., J Vis 2014). Metaphase preparations for the SCREEN and InSite assays followed the dGH protocol which includes UV- and exonuclease-mediated removal of the analog-incorporated daughter strand from metaphase chromosomes prior to probe hybridization (Ray et al., Chromosome 2013).

**K-Band Assay**

In this study, the K-Band assay encompassed culture of cryopreserved samples, staining and imaging of prepared slides, scoring and analysis of metaphase images, and generation of Genomic Integrity reports for each sample. Unlike conventional diagnostic G-banding, which is often used to assess constitutional or acquired cytogenetic abnormalities in a clinical context, KROMATID's K-Band assay is designed for genotoxicity assessment. Each cell is evaluated for the presence of structural variants, and results are aggregated to provide karyotype-level interpretation and quantitative variant frequency that enables detection of rare or low-frequency genomic alterations that can occur in engineered cell populations. Chromosome analysis follows the International System for Human Cytogenomic Nomenclature (ISCN) guidelines, and its recommendations cover the description of numerical and structural chromosomal changes detected using microscopic and cytogenetic techniques.

**Custom InSite Assay**

A custom InSite assay was developed to detect targeted integration of the CD38CAR transgene and to assess structural variation at the integration site. The assay employs three custom-designed probes: one targeting a 3.5 kb unique region of the CD38CAR sequence, labeled with ATTO-643, and two probes targeting approximately 2 Mb of genomic sequence immediately flanking the CD38 edit site on Chromosome 4 (999 kb telomeric and 958 kb centromeric). The telomeric flanking probe, labeled with ATTO-550, spans Chr 4 coordinates 14,809,769–15,768,219 (GRCh38), while the centromeric flanking probe, labeled with 6- FAM, spans Chr 4 coordinates 15,863,887–16,863,163. Probes were designed using a unidirectional tiled oligonucleotide strategy restricted to unique genomic sequences. Repetitive elements were masked, and probe specificity was validated via BLAST alignment against the human reference genome. Quality control testing for the flanking chromosome 4 probes was performed on metaphase spreads from unedited control samples to confirm accurate localization. The CD38CAR-specific probe was tested on edited samples known to contain the transgene to verify performance. Following qualification, the InSite assay was applied to two experimental samples: S028178 (WT γδT Cells) and S028182 (CD38KO/CD38CAR γδT Cells). Metaphase spreads were prepared and imaged using the same methods as for the SCREEN assay. Spreads were prequalified to ensure intact chromosome morphology and successful hybridization. Imaging was performed with appropriate excitation and emission filters for each fluorophore, allowing independent resolution of each probe signal. Analysis focused on probe localization and signal configuration to determine on-target versus o_-target integration and to detect structural rearrangements at the edit site. Each metaphase was assessed for chromosomal abnormalities at or near the CD38 integration locus, including translocations, truncations, inversions, sister chromatid exchanges, deletions, and insertions.

**Screen Assay**

For whole-genome analysis, KROMATID's SCREEN assay was hybridized to prepared metaphase spreads, counterstained with DAPI (Vectashield, Vector Laboratories, Newark, CA, USA), and imaged on an Applied Spectral Imaging Harmony system (Applied Spectral Imaging, Carlsbad, CA, USA) using a 100× objective. The assay consists of single-stranded, unidirectional tiled oligos designed to hybridize to unique sequences in each of the 24 human chromosomes. Each chromosome's oligos are end-labeled with one of five spectrally distinct fluorophores, combined such that chromosomes sharing the same color can still be differentiated by size, shape, and centromere position. Expected KROMASURE™ SCREEN signal patterns for the diploid human reference genome were used to construct preliminary karyograms for 50 cells (Bailey et al., J Pers Med 2024). These karyograms enabled perchromosome attribution of inter- and intra-chromosomal structural variation events, including inversions, translocations, aneuploidy (gain and loss), insertions, centromere abnormalities, and complex rearrangements for each sample. The assay detects translocations, insertions, and inversions at a much finer resolution than possible with KBand (~20 kb vs. 5-10 Mb), although it does not provide band-level attribution of breakpoint

### Supplementary Table 1. Reagents

| Reagent | Company | Catalog # |
|---|---|---|
| EasySep™ Human γδT Cell Isolation Kit | STEMCELL Technologies | 19225 |
| αPan-γδ TCR antibody (clone: 11F2 / REA591) | Miltenyi Biotec | 130-122-291 |
| In Vivo Ready™ Anti-Human CD28 (clone: CD28.2) | Tonbo Bioscience | 400289U100 |
| Human T Cell Media | Bio-Techne | CCM038-GMP-1L |
| CTS™ Immune Cell Serum Replacement | Gibco | A2596101 |
| 6-well G-Rex culture plates | Wilson Wolf | 80660M |
| CryoStor® CS10 | STEMCELL Technologies | 100-1061 |
| Directional Genomic Hybridization Cell Prep Kit | KROMATID | dGH 0001 |
| Colcemid block | KROMATID | COL 001 |
| Seahorse XF RPMI media | Aligent | 103576-100 |
| Calcein AM | Invitrogen | C3100MP |
| D-Luciferin, Sodium Salt (Proven and Published®) | GoldBio | LUCNA-1G |
| HiFi CRISPR-associated protein 9 (Cas9) Nuclease V3 | Integrated DNA Technologies | 1081061 |

### Supplementary Table 2. Flow Cytometry Antibodies

| Marker | Conjugate | Clone | Dilution (1 in X) | Company | Catalog # |
|---|---|---|---|---|---|
| αβTCR | PE | IP26 | 40 | Biolegend | 306708 |
| αβTCR | BV711 | IP26 | 40 | Biolegend | 306740 |
| CD3 | APC | UCHT1 | 40 | Biolegend | 300412 |
| CD3 | BV421 | UCHT1 | 40 | Biolegend | 300434 |
| CD38 | APC | HIT2 | 40 | Biolegend | 303510 |
| CD56 | PE-Vio615 | REA196 / NCAM16.2 | 40 | Miltenyi Biotec | 130-114-550 |
| Vδ1 | FITC | TS8.2 | 40 | Invitrogen | TCR2730 |
| Vδ2 | PE/Cy7 | B6 | 40 | Biolegend | 331421 |
| Whitlow 218 Linker | PE | E3U7Q | 40 | Cell Signaling | 62405S |
| γδ TCR | BV421 | B1 | 20 | BD Horizon | 562560 |
| Viability | Zombie NIR | | 200 | Biolegend | 423105 |
| Glut1 | FITC | 202915 | 5 | R&D Systems | FAB1418F |
| CD71 | BV711 | M-A712 | 10 | BD Biosciences | 563767 |
| CD98 | PE | UM7F8 | 50 | BD Biosciences | 556077 |

**Supplementary Table 3. CyTOF Antibodies**

| Metal | Antibody | Clone | Vendor | Catalog # | Concentration |
|---|---|---|---|---|---|
| 89 | CD235 | HIR2 | Biolegend | | 2 ug/ml |
| 111 | Lysozyme | BGN/0696/5B1 | Invitrogen | MA1-82873 | 2 ug/ml |
| 112 | GAPDH | AM4300 | Invitrogen | AM4300 | 2 ug/ml |
| 113 | CD3 | UCHT1 | Biolegend | 300402 | 2 ug/ml |
| 115 | CD45 | HI30 | Biolegend | 304002 | 2 ug/ml |
| 116 | H3K9ac | C5B11 | CST | 96075SF | 2 ug/ml |
| 139 | CD41 | HIP8 | Biolegend | 303702 | 2 ug/ml |
| 140 | cPARP | F21-A52 | BD | 51-9000017 | 2 ug/ml |
| 141 | CD27 | M-T271 | Biolegend | 356401 | 1 ug/ml |
| 142 | CD7 | M-T701 | BD | 555359 | 2 ug/ml |
| 143 | CD71 | CY1GG4 | Biolegend | 334102 | 2 ug/ml |
| 144 | CD94 | HP-3D9 | BD | 555887 | 2 ug/ml |
| 145 | CD4 | RPA-T4 | Biolegend | 300502 | 2 ug/ml |
| 146 | CD8a | RPA-T8 | Biolegend | 301002 | 2 ug/ml |
| 147 | CD56 | NCAM16.2 | BD | 559043 | 1 ug/ml |
| 148 | CD34 | 581 | Biolegend | 343502 | 2 ug/ml |
| 149 | HLAE | 3D12 | Biolegend | 342602 | 4 µg/ml |
| 150 | CD117 | 104D2 | Biolegend | 313202 | 2 ug/ml |
| 151 | CD123 | 6H6 | Biolegend | 306002 | 2 ug/ml |
| 152 | CD33p67 | p97.6 | Biolegend | 366602 | 2 ug/ml |
| 153 | HLA-DR | L243 | Biolegend | 307602 | 2 ug/ml |
| 154 | CD69 | FN50 | Biolegend | 310902 | 2 ug/ml |
| 155 | TCR a/B | IP26 | Biolegend | 306702 | 4 µg/ml |
| 156 | CD11c | S-HCL-3 | Biolegend | 371502 | 1 µg/ml |
| 157 | CD45RA | HI100 | Biolegend | 304102 | 1 µg/ml |
| 158 | Ki-67 | SolA15 | Invitrogen | 14-5698-82 | 4 µg/ml |
| 159 | CD38 | HIT2 | Biolegend | 303502 | 2 ug/ml |
| 160 | CD14 | M5E2 | Biolegend | 301802 | 3 ug/ml |
| 161 | CD16 | 3G8 | Biolegend | 302002 | 2 ug/ml |
| 162 | CD11b | ICRF44 | Biolegend | 301302 | 3 ug/ml |
| 163 | CD64 | 10.1 | Biolegend | 305002 | 2 ug/ml |
| 164 | CD15 | W6D3 | Biolegend | 323002 | 2 ug/ml |
| 165 | pRb | J112-906 | BD | 558389 | 1.5 µg/ml |
| 166 | NKG2D/CD314 | 1D11 | Biolegend | 320802 | 4 µg/ml |
| 167 | CD99 | HCD99 | Biolegend | 371302 | 2 ug/ml |
| 168 | CD13 | WM15 | Biolegend | 301702 | 2 ug/ml |
| 169 | p16 | D71M | CST | 50604SF | 4 µg/ml |
| 170 | CD107a | H4A3 | Biolegend | 328602 | 2 ug/ml |
| 171 | TCR Vg1 | REA173 | Miltenyi Biotec | 130-122-285 | 2 ug/ml |
| 172 | TCR Vg2 | b6 | Biolegend | 331402 | 2 ug/ml |
| 173 | CD19 | HIB19 | Biolegend | 302202 | 2 ug/ml |
| 174 | CD20 | 2H7 | Biolegend | 302302 | 4 µg/ml |
| 175 | Perforin | dG9 | Biolegend | 308102 | 4 µg/ml |
| 176 | pCREB | 87G3 | CST | 39561SF | 2 µg/ml |
| 194 | H3K27me3 | C36B11 | CST | 9733BF | 3 µg/ml |
| 196 | Vamp7 | 549115 | R&D | MAB6117 | 3 µg/ml |
| 198 | Total Histone H1 | AE-4 | Bio-Rad | 4974-7808 | 2 µg/ml |
| 209 | pHH3 | HTA28 | Biolegend | 641002 | 1 ug/ml |

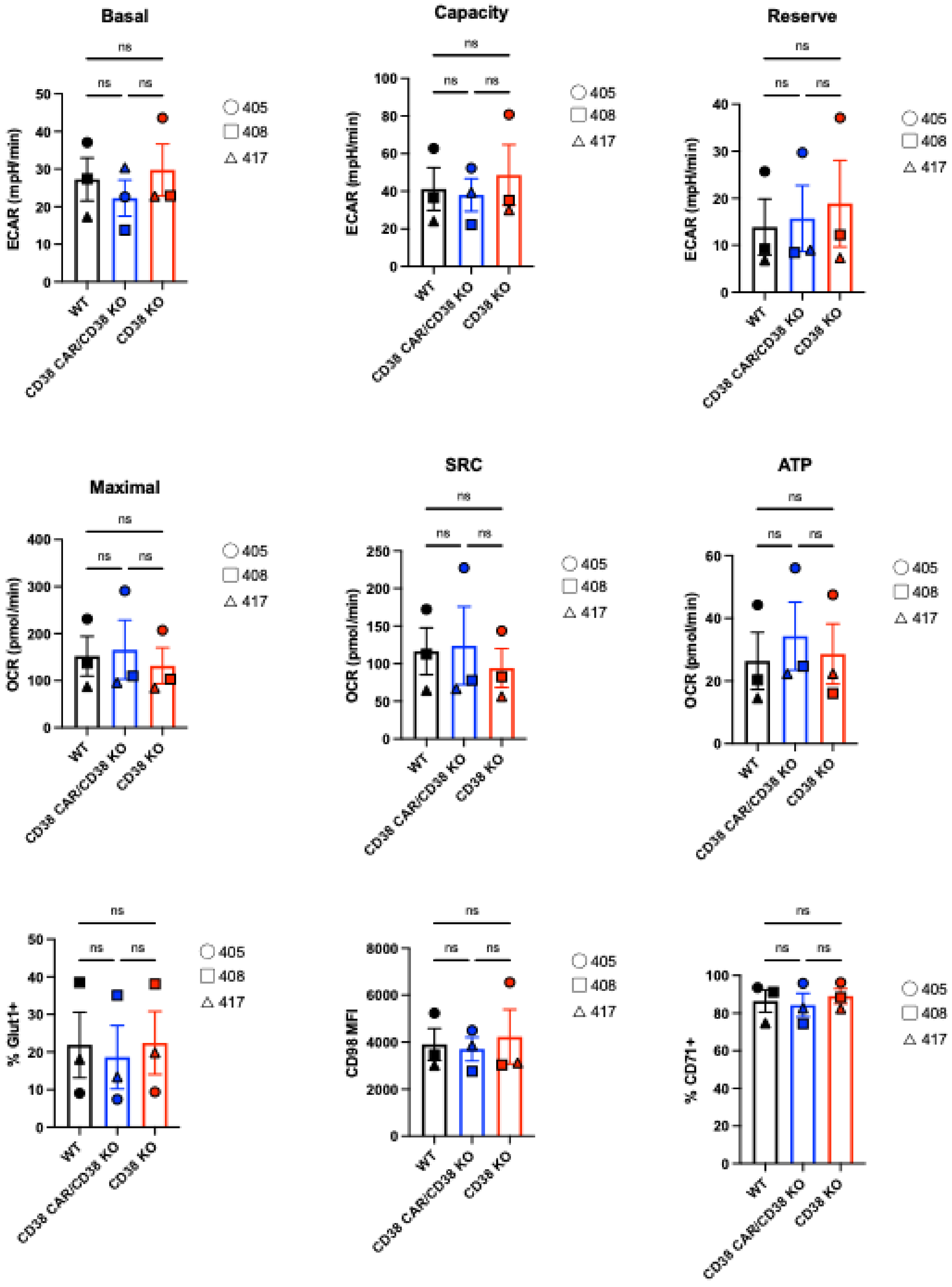


**Supplementary Figure 1.** Metabolic assays showed no significant changes in γδT cells post editing and CD38 deletion.

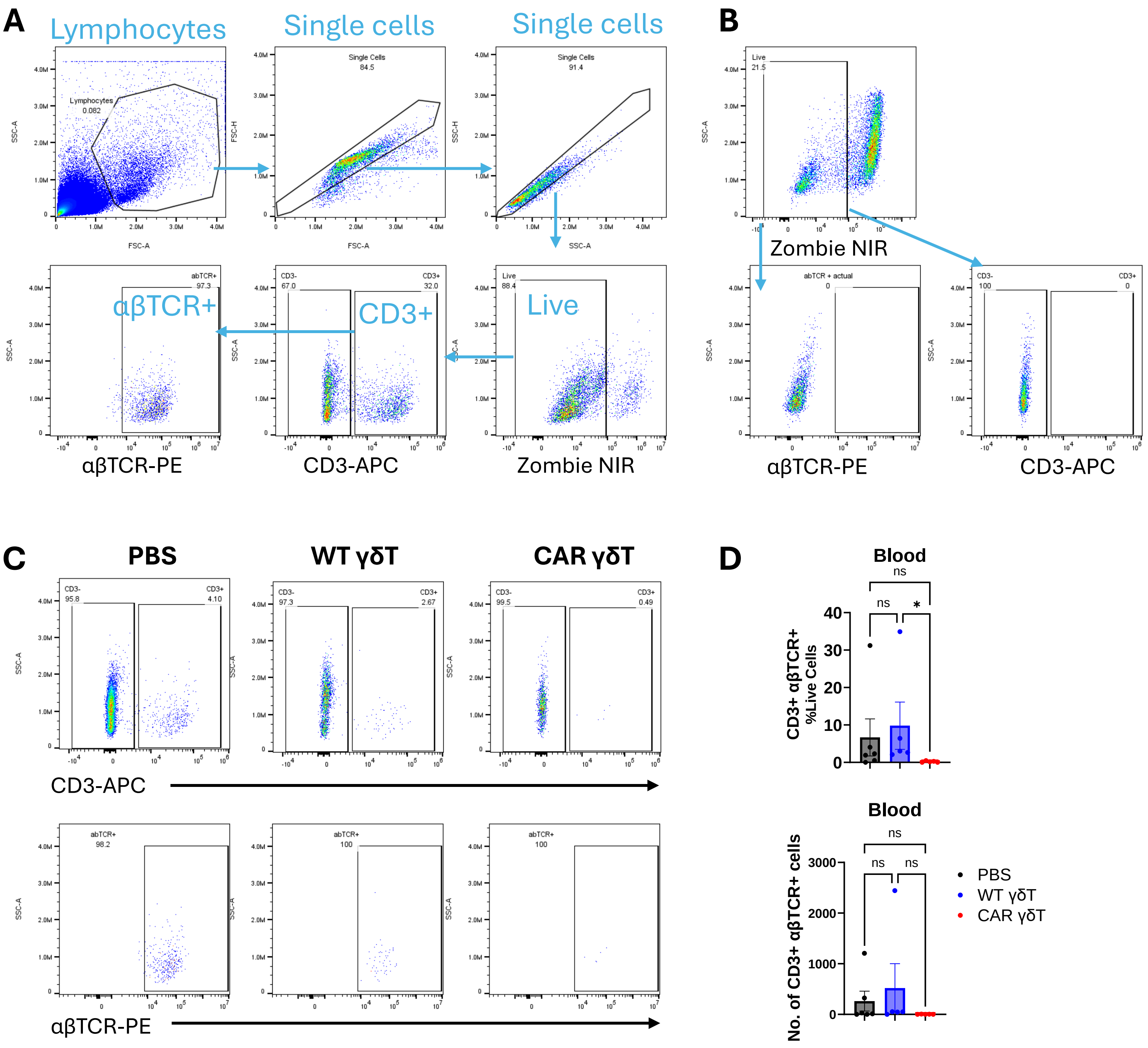


**Supplemental Figure 2**. Flow cytometry analysis of blood collected from T-ALL bearing mice. NSG mice engrafted with Luc$^{+}$ Jurkat cells were treated with PBS, WT γδT, or CD38KO/CD38CAR γδT cells (n≥5 per cohort). Blood was collected on autopsy, stained, and analyzed by flow cytometry for residual T-ALL cells. (A) Gating strategy. (B) Live/dead single stain control for marker gating (CD3 and αβTCR). (C) Representative flow plots of CD3+ and αβTCR+ cells from blood samples of PBS, WT γδT, and CAR γδT treated mice. (D) Graphs quantifying CD3+ αβTCR+ proportion of total live cells (top) or total number of CD3+ αβTCR+ cells (bottom). Each shape represents one animal. Data represents mean ± SEM. Statistical analysis performed using Kruskal-Wallis non-parametric test with Dunn's multiple comparison correction.
